# Factors Related to Dental Caries, Periodontal Disease, and Psychological effect of those diseases: A Structural Equation Modeling and Artificial Neural Network model comparative approach

**DOI:** 10.1101/364539

**Authors:** Capt Sirsendu Ghosh

## Abstract

Exceptional growth in the development of oral health of various populations worldwide over the last three decades cannot lessen tribulations in dental caries, periodontal disease, and psychological problems, which are still prevalent in many communities, especially among the poor socioeconomic groups in developing countries like India. Dental caries and periodontal disease are exceedingly related to the lifestyle associated risk factors and various daily habits including smoking and tobacco chewing. Dietary habit is one of the prime causative behind the formation of dental caries and simultaneously the dietary habit is greatly influenced by the person’s socio-economic status. In this study to explore the factors related to dental caries and periodontal disease and how these diseases manipulate the mental health of the people, some SEMs and some ANN models are also formed. At last both models are compared and explained about their purposes and usability for further applications.

## Background

Exceptional growth in development of oral health of various populations worldwide over the last three decades cannot lessen tribulations in dental caries, periodontal disease and psychological problems, which are still prevalent in many communities, especially among the poor socioeconomic groups in developing countries [1] like India. Dental caries and periodontal disease are exceedingly related to the lifestyle associated risk factors and various daily habits including smoking and tobacco chewing. Dietary habit is one of the prime causative behind the formation of dental caries and simultaneously the dietary habit is greatly influenced by the person’s socio-economic status. Based on the previous studies and from the stand point of public health, in this study socio-economic factor, oral health knowledge, dietary habits, tobacco using habit, psychological factor related to caries and periodontal disease was studied. The KAP [Knowledge, Attitude, and Practice] model has huge influence in this study. Although many factors associated to dental caries and periodontal disease have been recognized in many previous studies using multiple regression [4, 5] models and structural equation models [22], but still more advanced technique is required to establish the relation among the disease causing factors and diseases. The multiple regression models are not much successful to distinguish between correlation and causal associations present between the disease and factor. Similarly the structural equation modeling approach has some assumptions regarding the linear relation between factors and diseases, which is not true in all real life scenarios. From the prediction point of view, it’s a matter of judgment that which is better, a structural equation model or an artificial neural network model. So there is a need to apply such kind of procedure which can capture the nonlinear relationship among the factors and diseases, side by side will be able to predict the consequence with some accuracy. Previously structural equation modeling [7] based study was done on Chinese children population [22] but no such complex models are tested on general population of India. Intension of this study is to employ the structural equation modeling and artificial neural network model concurrently on the data, which covers diverse age groups of the population. On the basis of oral heath knowledge, attitude and practice [7], few hypothesizes are tested by using different Structural Equation Method (SEM) and Artificial Neural Network (ANN) models. It was hypothesized that socio-economic status has an impact on the dental caries and periodontal diseases. Oral hygiene practice or maintenance and dietary habit has an impact on carious condition of teeth including total life time caries experience, which includes total number of decayed, missing and filled teeth present in participants’ oral cavity. Tobacco smoking and using other kind of smokeless tobacco products greatly influence the formation of periodontal disease. Caries and it’s after effect like filled or missing teeth and periodontal diseases creates a psychological impact on participant’s mind. Missing teeth due to caries, tobacco usage, dietary habit, periodontal disease etc. combined creates an impact on the psychological state of the person. In this study to explore the factors related to dental caries and periodontal disease and how these diseases manipulate the mental health of the people, some SEMs and some ANN models are also formed. At last both models are compared and explained about their purposes and usability for further applications.

## Methods

### Study subjects

A cross sectional study was conducted from June to September 2015 in the six villages of Ballabhgarh district, Haryana, India with a target sample size of 960. Ultimately 829 people participated in this study, giving a response rate of 86.35%. Four age groups are included in this study 5 – 7 years, 12 – 15 years, 35 – 44 years, 65 – 74 years age of people. A probabilistic approach was maintained during the sampling of the study subjects. Prior to the study, ethical approval from the ethics committee of the All India Institute of Medical Sciences, New Delhi and written consent from every participant’s were obtained. In case of 5-7 years, 12-15 years age group of people, the consent was obtained from their parents.

### Questionnaire

A questionnaire study [10] was conducted on the study subjects. Factors included in questionnaire were: Socio-economic status of participant [Table 1a], participant’s oral heath knowledge [Table 1b], dietary habits [Table 1c], tobacco use habit [Table 1g] (5-7 years, 12-15 years were exempted from these questions), psychological status [Table. 1d], oral hygiene practice or maintenance [Table 1f] and knowledge [Table 1e]. SES was measured by participant’s education, participant’s father’s education, participant’s mother’s education, participant’s occupation, participant’s father’s occupation, participant’s mother’s occupation [participant’s father’s and mother’s education and occupation were asked only in case of 5-7 years, 12-15 years age group subjects]. Participant’s oral health knowledge was measured by participant’s awareness about: number of natural teeth present in his/her mouth and state of those teeth according to him/her. Dietary habit was measured by consumption history of following foods: Fresh fruits, Biscuits/cakes/buns/gur/sweet pies, and Soft drinks/lemonade/coca cola. Tobacco using habit was measured by the questions: current smoker, current user of smokeless tobacco, previous smoker, and previous user of smokeless tobacco. Oral hygiene maintenance practice and knowledge were measured by: Frequency of teeth cleaning, tooth paste user or not, whether that tooth paste contained fluoride or not, uses toothbrush, uses wooden and plastic toothpick, uses datum for teeth cleaning, cleans teeth by finger, uses threads for teeth cleaning, uses coal or any other cleaning aid to clean his/her teeth. Psychological status was measured by the following questions: Felt tense due to oral problems, Embarrassed due to appearance of teeth, Avoid smiling due to teeth, Sleep often interrupted due to oral problems, Days not at work due to oral problems, Difficulty in usual activity due to oral condition, Less tolerant to spouse or people nearer to him/her due to oral condition, Reduced participation in social activities due to oral problems, Embarrassed due to bad breath. Frequency distribution and ordinal levels are presented at [Table no.1 a to g].

**Table No. 1a:**
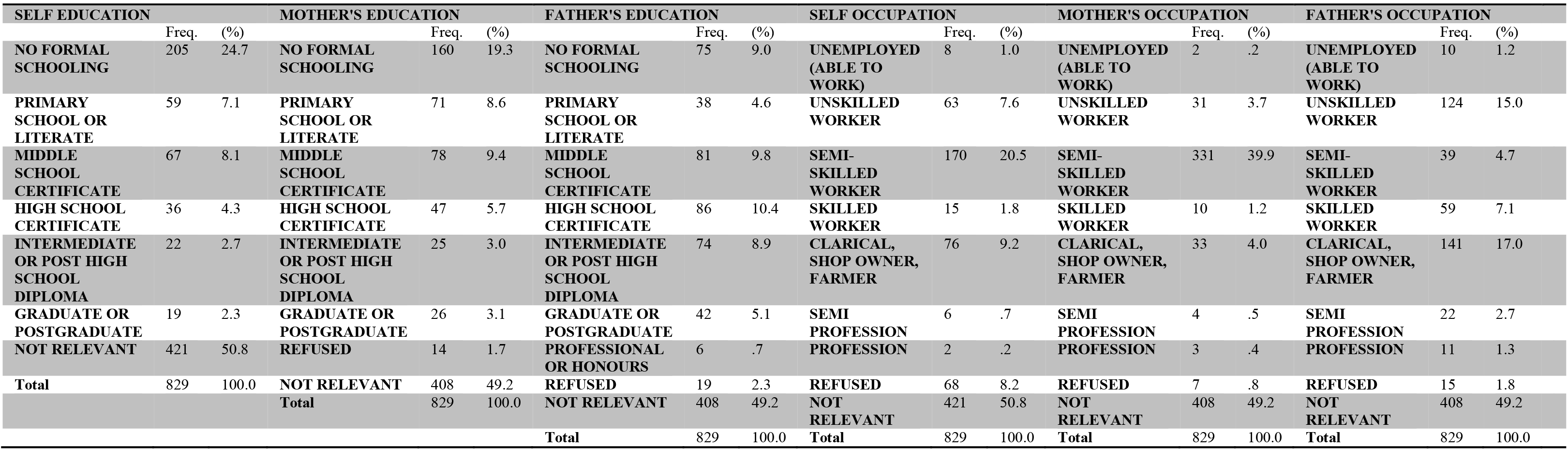
Socio-economic factor.

**Table No. 1b:**
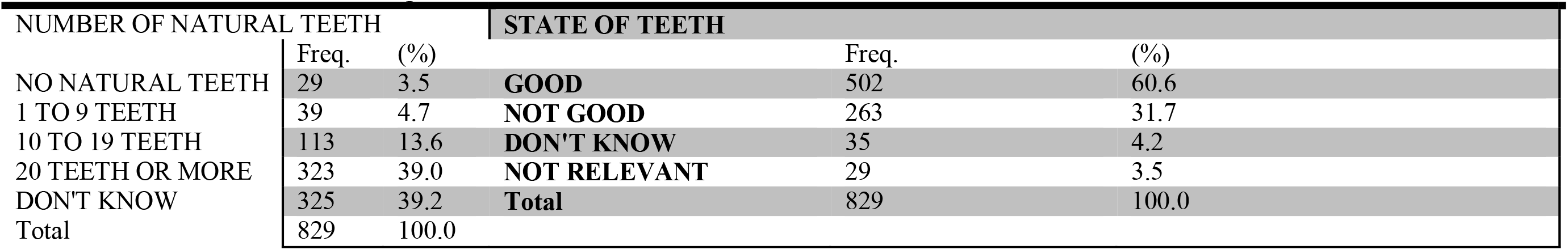
Oral health knowledge/Awareness.

| NUMBER OF NATURAL TEETH |  |  | STATE OF TEETH |  |  |
| --- | --- | --- | --- | --- | --- |
|  | Freq. | (%) |  | Freq. | (%) |
| NO NATURAL TEETH | 29 | 3.5 | GOOD | 502 | 60.6 |
| 1 TO 9 TEETH | 39 | 4.7 | NOT GOOD | 263 | 31.7 |
| 10 TO 19 TEETH | 113 | 13.6 | DON'T KNOW | 35 | 4.2 |
| 20 TEETH OR MORE | 323 | 39.0 | NOT RELEVANT | 29 | 3.5 |
| DON'T KNOW | 325 | 39.2 | Total | 829 | 100.0 |
| Total | 829 | 100.0 |  |  |  |

**Table No. 1c:**
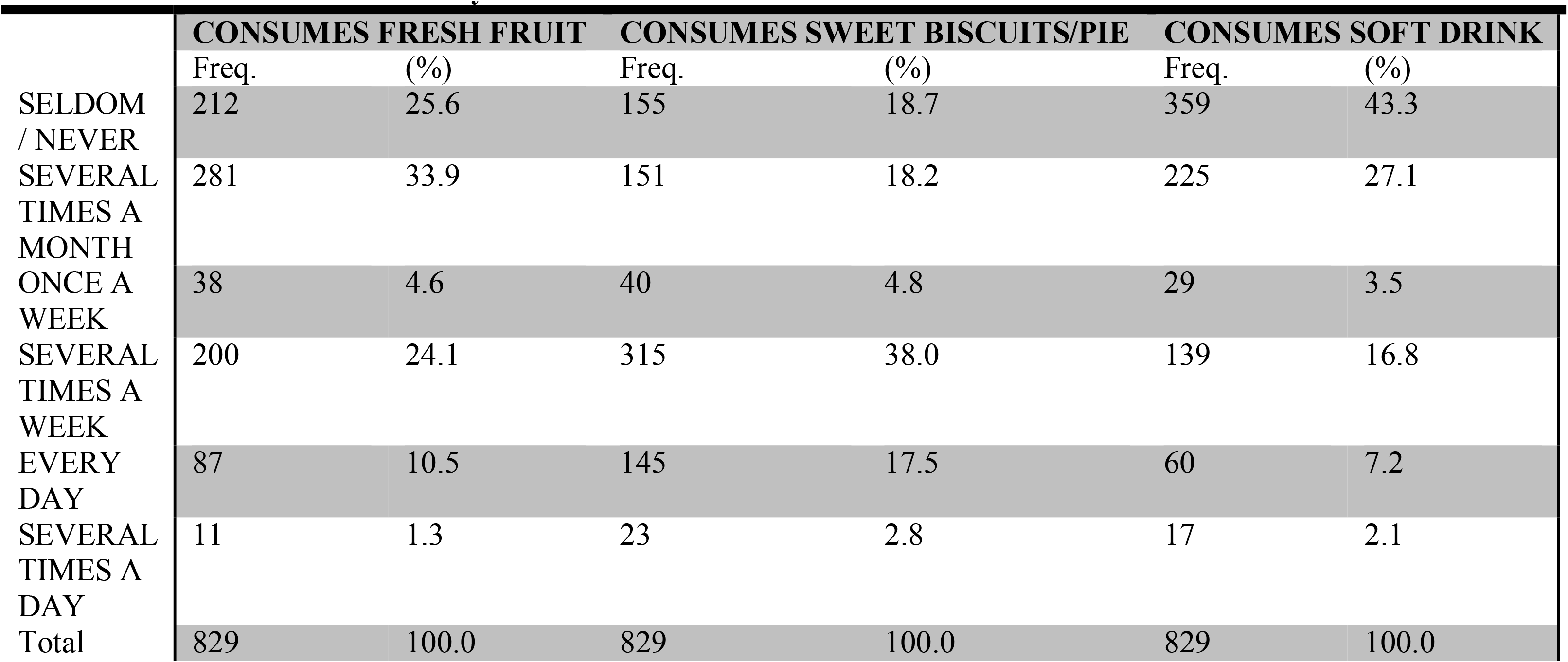
Dietary habit factor.

**Table 1d:** Psychological factor.

| FELT TENSE DUE TO ORAL PROBLEMS |  |  | EMBARRASSED DUE TO APPEARANCE OF TEETH |  |  | AVOID SMILING DUE TO TEETH |  |  | INTERUPTTED SLEEP DUE TO ORAL PROBLEM |  |  |
| --- | --- | --- | --- | --- | --- | --- | --- | --- | --- | --- | --- |
|  | Freq. | (%) |  | Freq. | (%) |  | Freq. | (%) |  | Freq. | (%) |
| YES | 29 | 3.5 | YES | 16 | 1.9 | YES | 11 | 1.3 | YES | 34 | 4.1 |
| NO | 800 | 96.5 | NO | 813 | 98.1 | NO | 818 | 98.7 | NO | 795 | 95.9 |
| Total | 829 | 100.0 | Total | 829 | 100.0 | Total | 829 | 100.0 | Total | 829 | 100.0 |
| NOT AT WORK DUE TO ORAL PROBLEM |  |  | DIFFICULTY IN USUAL ACTIVITIES DUE TO ORAL PROBLEM |  |  | LESS TOLERANT TO SPOUSE/NEARER PERSON |  |  | REDUCED PARTICIPATION IN SOCIAL ACTIVITY |  |  |
|  | Freq. | (%) |  | Freq. | (%) |  | Freq. | (%) |  | Freq. | (%) |
| YES | 13 | 1.6 | YES | 7 | .8 | YES | 6 | .7 | YES | 5 | .6 |
| NO | 816 | 98.4 | NO | 822 | 99.2 | NO | 823 | 99.3 | NO | 824 | 99.4 |
| Total | 829 | 100.0 | Total | 829 | 100.0 | Total | 829 | 100.0 | Total | 829 | 100.0 |
| EMBARRASSED DUE TO BAD BREATH |  |  |  |  |  |  |  |  |  |  |  |
|  | Freq. | (%) |  |  |  |  |  |  |  |  |  |
| YES | 89 | 10.7 |  |  |  |  |  |  |  |  |  |
| NO | 740 | 89.3 |  |  |  |  |  |  |  |  |  |
| Total | 829 | 100.0 |  |  |  |  |  |  |  |  |  |

**Table No. 1e:**
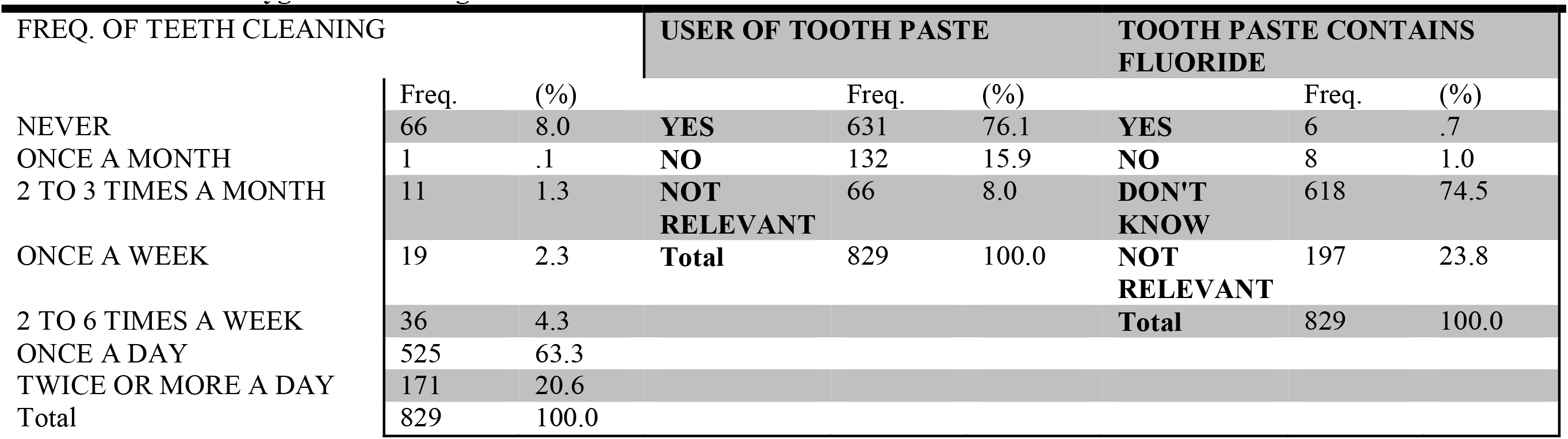
Oral hygiene Knowledge.

| FREQ. OF TEETH CLEANING |  |  | USER OF TOOTH PASTE |  |  | TOOTH PASTE CONTAINS FLUORIDE |  |  |
| --- | --- | --- | --- | --- | --- | --- | --- | --- |
|  | Freq. | (%) |  | Freq. | (%) |  | Freq. | (%) |
| NEVER | 66 | 8.0 | YES | 631 | 76.1 | YES | 6 | .7 |
| ONCE A MONTH | 1 | .1 | NO | 132 | 15.9 | NO | 8 | 1.0 |
| 2 TO 3 TIMES A MONTH | 11 | 1.3 | NOT RELEVANT | 66 | 8.0 | DON'T KNOW | 618 | 74.5 |
| ONCE A WEEK | 19 | 2.3 | Total | 829 | 100.0 | NOT RELEVANT | 197 | 23.8 |
| 2 TO 6 TIMES A WEEK | 36 | 4.3 |  |  |  | Total | 829 | 100.0 |
| ONCE A DAY | 525 | 63.3 |  |  |  |  |  |  |
| TWICE OR MORE A DAY | 171 | 20.6 |  |  |  |  |  |  |
| Total | 829 | 100.0 |  |  |  |  |  |  |

**Table No. 1f:** Oral hygiene maintenance practice/habit.

|  | **USER OF TOOTHBRUSH** |  | **USER OF WOODEN TOOTHPICK** |  | **CLEANS TEETH BY THREAD** |  | **USER OF DATUN** |  |
| --- | --- | --- | --- | --- | --- | --- | --- | --- |
|  | Freq. | (%) | Freq. | (%) | Freq. | (%) | Freq. | (%) |
| YES | 630 | 76.0 | 84 | 10.1 | 3 | .4 | 226 | 27.3 |
| NO | 133 | 16.0 | 679 | 81.9 | 760 | 91.7 | 537 | 64.8 |
| NOT RELEVANT | 66 | 8.0 | 66 | 8.0 | 66 | 8.0 | 66 | 8.0 |
| Total | 829 | 100.0 | 829 | 100.0 | 829 | 100.0 | 829 | 100.0 |
|  | **CLEANS TEETH BY FINGER** |  | **USER OF PLASTIC TOOTHPICK** |  | **CLEANS TEETH BY COAL** |  | **USES OTHER CLEANING AID** |  |
|  | Freq. | (%) | Freq. | (%) | Freq. | (%) | Freq. | (%) |
| YES | 266 | 32.1 | 7 | .8 | 15 | 1.8 | 58 | 7.0 |
| NO | 497 | 60.0 | 756 | 91.2 | 748 | 90.2 | 705 | 85.0 |
| NOT RELEVANT | 66 | 8.0 | 66 | 8.0 | 66 | 8.0 | 66 | 8.0 |
| Total | 829 | 100.0 | 829 | 100.0 | 829 | 100.0 | 829 | 100.0 |

**Table No. 1g:** Tobacco using habit.

|  | **CURRENT DAILY SMOKER** |  | **PAST DAILY SMOKER** |  | **CURRENT DAILY SMOKELESS TOBACCO USER** |  | **PAST DAILY SMOKELESS TOBACCO USER** |  |
| --- | --- | --- | --- | --- | --- | --- | --- | --- |
|  | Freq. | (%) | Freq. | (%) | Freq. | (%) | Freq. | (%) |
| YES | 144 | 17.4 | 48 | 5.8 | 36 | 4.3 | 18 | 2.2 |
| NO | 478 | 57.7 | 438 | 52.8 | 585 | 70.6 | 581 | 70.1 |
| NOT RELEVANT | 207 | 25.0 | 343 | 41.4 | 208 | 25.1 | 230 | 27.7 |
| Total | 829 | 100.0 | 829 | 100.0 | 829 | 100.0 | 829 | 100.0 |

### Clinical Examination

Oral examinations were conducted by dental surgeons. Record of number of decayed, missing, filled teeth due to caries, community periodontal index [CPI], and loss of periodontal attachment [LOA] were based on the criteria recommended by world health organization [WHO] [10]. In case of 5-7 years age group subjects only dmft was measured. In case of 12-15 years age group subjects’ DMFT, CPI was measured. In case of 35-44 years, 65-74 years age group subjects DMFT, CPI, LOA were measured. Numbers of decayed, missing, filled teeth were measured as continuous variable. CPI and LOA were measured as ordinal variable. [Table 2a and 2b]

**Table No. 2a:**
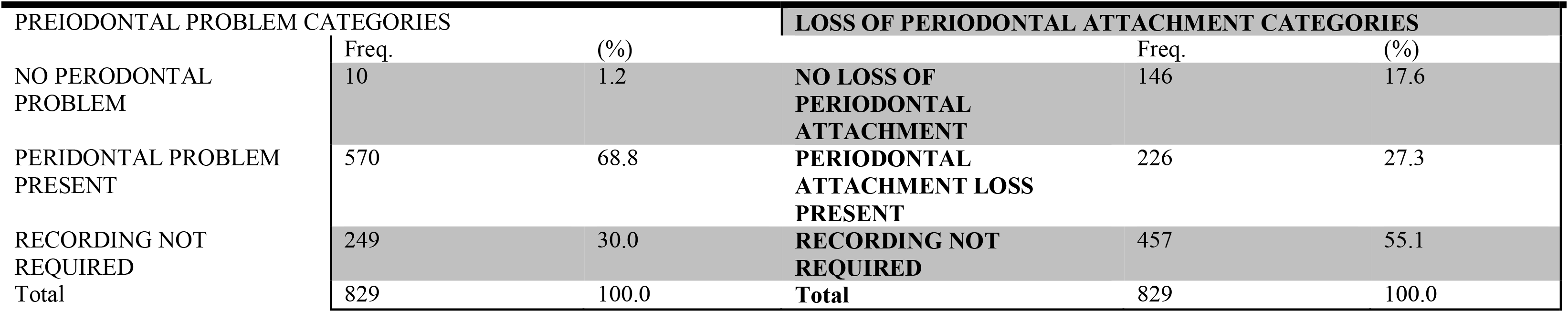
Periodontal disease.

**Table No. 2b:** Decayed, Missing, Filled Teeth.

|  | N | Mean | Std. Deviation | Variance |
| --- | --- | --- | --- | --- |
| Caries calculated | 829 | 1.0748 | 1.69090 | 2.859 |
| Missing calculated | 829 | .8179 | 2.09770 | 4.400 |
| Filled calculated | 829 | .0796 | .44606 | .199 |

### Data Analysis

For data analysis and artificial neural network modeling SPSS for windows version 19.0 and for structural equation modeling STATA version 12 were used. At first different hidden constructs or factors were determined by conducting Categorical Principal Component analysis [CATPCA]. According to factor loadings, different factors were decided [Table No. 3]. Observations under each variable were standardized before creating the structural equation model. Kernel density plots were checked before and after standardization and those plots were super imposed by normal distribution curve. There were no changes in Kernel density plot after standardization of variables. [Supplementary Figure 19]

**Table 3:** Loading Factors.

| Factors | Constructing Variables | Levels of response of each variables | Type of variable | Loadings Respectively |
| --- | --- | --- | --- | --- |
| Oral Hygiene Measure | Freq. of teeth cleaning*, Usage of tooth paste, Tooth paste containing fluoride**, Use tooth brush, Use wooden toothpick, Use Finger to clean teeth, Use plastic toothpick, Use threads, Use charcoal to clean teeth, Use Datun, Use other cleaning aid | * 8 levels, **4 levels, all others 3 levels | Ordinal | -0.972, 0.976, 0.605, 0.976, 0.972, 0.972, 0.972, 0.972, 0.972, 0.972, 0.972 |
| Specific Diet | Consumption of Fresh fruit, Biscuits/cakes/buns/gur/sweet pies, Soft drinks/lemonade/Coca-cola | 6 levels | Ordinal | 0.614, 0.517, 0.606 |
| Socioeconomic Status | #Father's Education*, #Mother's Education*, Self(Subject's) education*, #Father's occupation**, #Mother's occupation**, Self(Subject's) occupation** | *8 levels, **10 levels, # are only applicable in case of 12-15 yrs & 5-7 yrs age group's subjects | Ordinal | 0.930, 0.930, -0.930, 0.930, 0.931, -0.930 |
| Tobacco using habit | Current smoker, Past smoker, Currently use smokeless tobacco, In past used smokeless tobacco | 3 levels | Ordinal | -0.772, 0.754, -0.773, -0.710 |
| Psychological effect | Felt tense due to oral problems, Embarrassed due to appearance of teeth, Avoid smiling due to teeth, Sleep often interrupted due to oral problems, Days not at work due to oral problems, Difficulty in usual activity, Less tolerant of spouse or people close to you, Reduced participation in social activities, Embarrassed due to bad breath | 3 levels | Ordinal | 0.393, 0.646, 0.559, 0.528, 0.652, 0.676, 0.601, 0.563, 0.417 |
| Caries effect | Total carious teeth of each subject, Total missing teeth of each subject, Total Filled teeth of each subject | None | Continuous Data | 0.822, 0.598, 0.672 |

According to previously planned hypothesizes six different structural equation models were Constructed and their validity were tested. The observed variables were defined as formative items of their constructs in the measurement model. Since some observed variables were in continuous and some others in ordinal, asymptotic distribution free estimation method was used in each structural equation model [SEM]. The magnitude and significance of the hypothesized links were evaluated with the β [β = path coefficient] and P-values. A P-value less than 0.05 were considered as statistically significant. The comparative fit index [CFI], the Tuker-Lewis index [TLI], the Root means square error approximation [RMSEA] were used to measure the goodness of fit of the model. CFI and TLI value ≥ 0.90, P-close > 0.05, RMSEA ≤ 0.06 served as criteria of model fitness [11, 12, 22]. A model was accepted when all the criteria’s were fulfilled; if any one criterion remains unfulfilled then the model was rejected. As the distribution of many variables were non-normal with high skewness and kurtosis so, chi-square was not considered as a measure of goodness of fit [Table No. 7]. In case of non normal data the comparative fit measure were less affected. CFI was not computed if the RMSEA of null model was less than 0.158. Neural network models were formed by using the non standardized variables [Table No. 4]. The variables which were included in structural equation models were considered during formation of artificial neural network model [ANN]. Multilayer perceptron model with a partition of Training: Testing: Holdout = 7:4:1 was considered in each case of model. All models were trained online. In case of continuous dependant variables sum of square error and average overall relative error (in %) were measured. In case of categorical dependant variable cross entropy error and average (%) incorrect prediction were measured. Activation functions for hidden layers were hyperbolic tangent in each case. Activation function for categorical dependant variables were softmax and activation function for continuous dependant variables were identity. Continuous input/ independent variables were considered as covariates [Table No. 5 and 6].

**Table No. 4:**
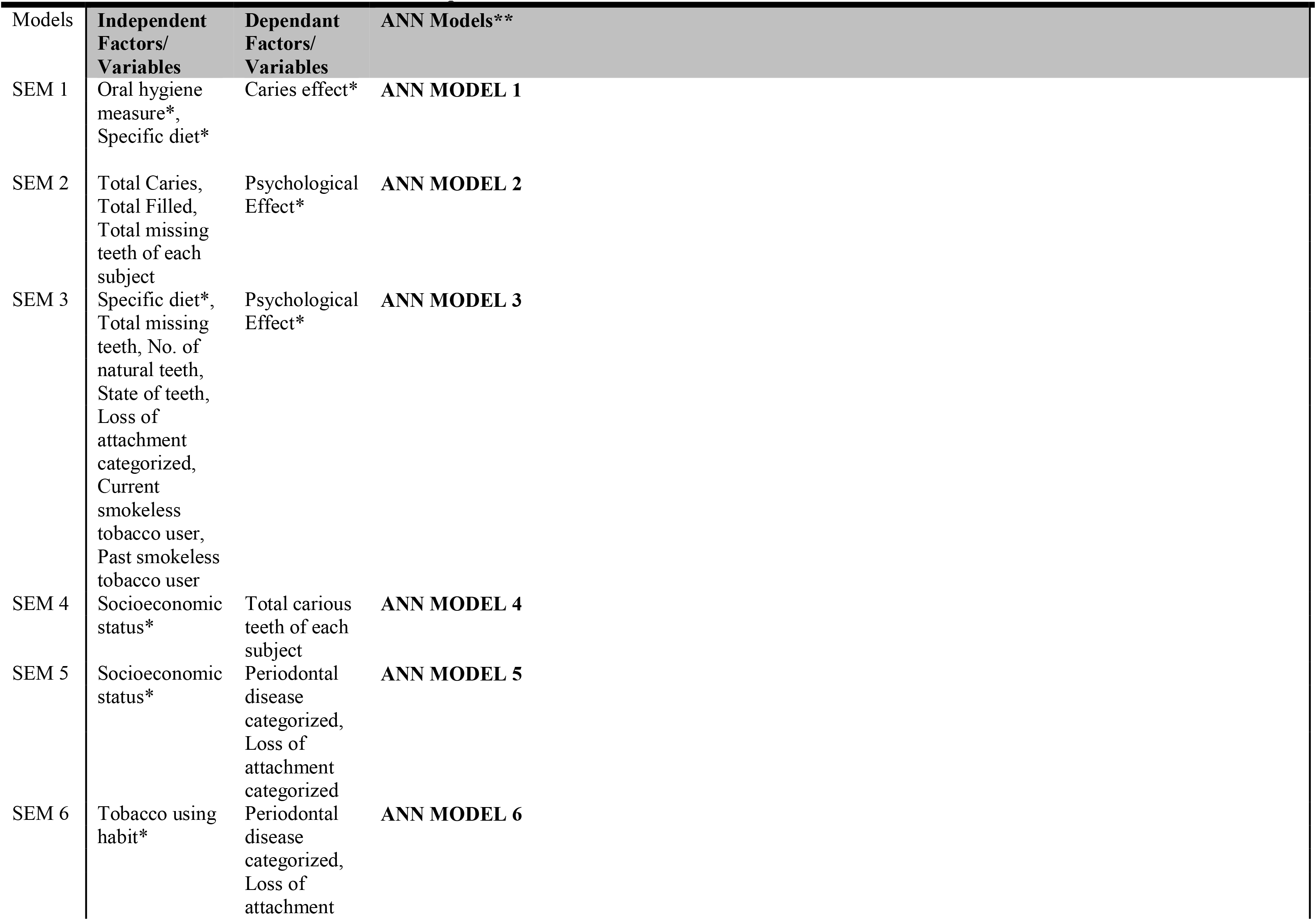
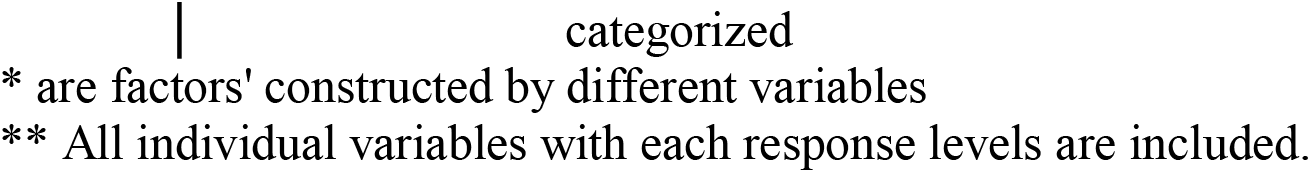
Constructional units of Structural Equation Models and Artificial Neural Networks.

| Models | Independent Factors/<br>Variables | Dependant Factors/<br>Variables | ANN Models** |
| --- | --- | --- | --- |
| SEM 1 | Oral hygiene measure*,<br>Specific diet* | Caries effect* | **ANN MODEL 1** |
| SEM 2 | Total Caries,<br>Total Filled,<br>Total missing teeth of each subject | Psychological Effect* | **ANN MODEL 2** |
| SEM 3 | Specific diet*,<br>Total missing teeth, No. of natural teeth, State of teeth, Loss of attachment categorized, Current smokeless tobacco user, Past smokeless tobacco user | Psychological Effect* | **ANN MODEL 3** |
| SEM 4 | Socioeconomic status* | Total carious teeth of each subject | **ANN MODEL 4** |
| SEM 5 | Socioeconomic status* | Periodontal disease categorized, Loss of attachment categorized | **ANN MODEL 5** |
| SEM 6 | Tobacco using habit* | Periodontal disease categorized, Loss of attachment | **ANN MODEL 6** |

\* are factors' constructed by different variables
\*\* All individual variables with each response levels are included.

**Table No. 5:**
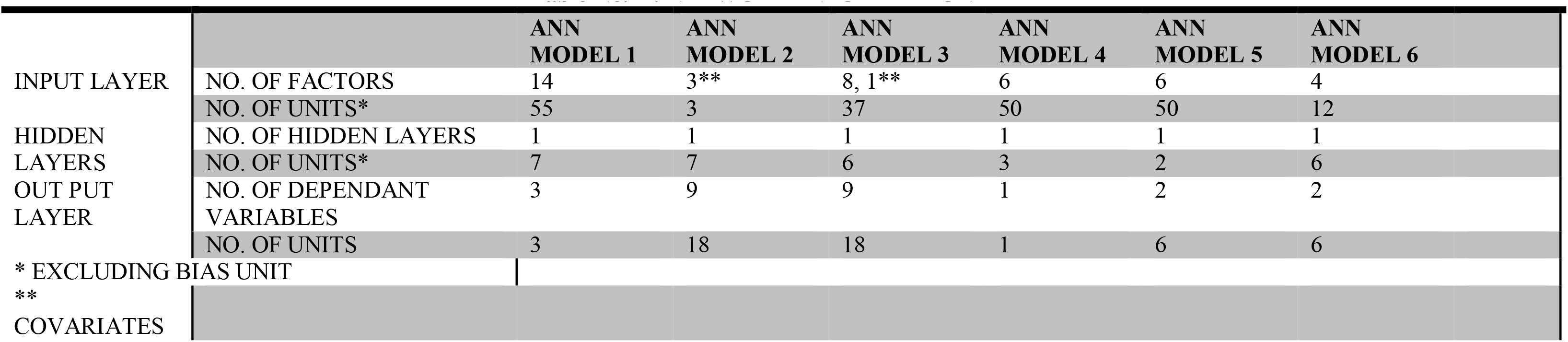
NETWORK INFORMATION.

**Table No. 6:**
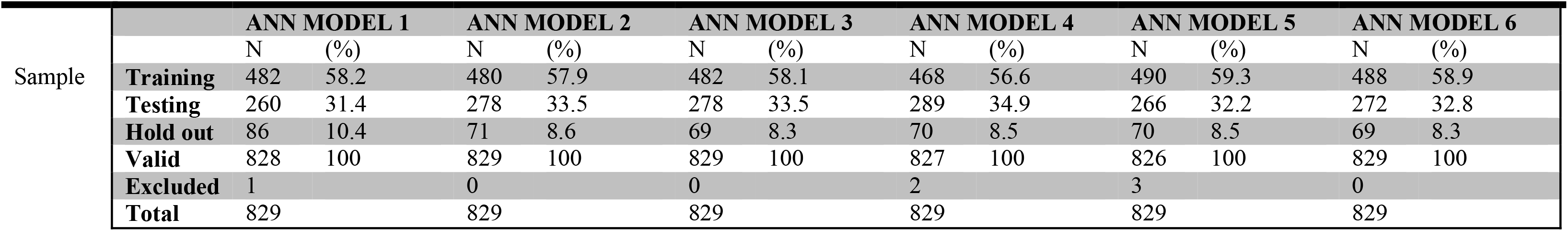
MULTILAYER PERCEPTRON NEURAL NETWORK MODEL:- CASE PROCESSING SUMMARY.

**Table No. 7:-.**
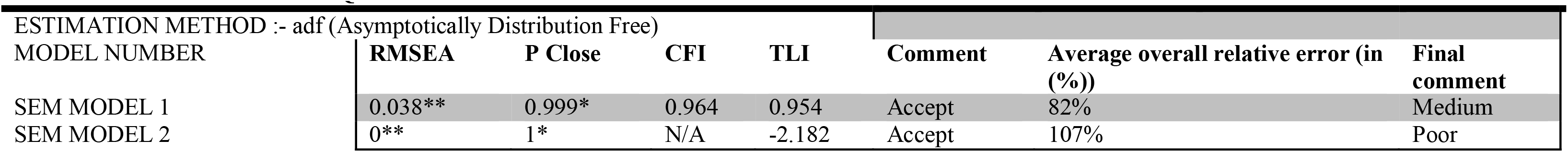
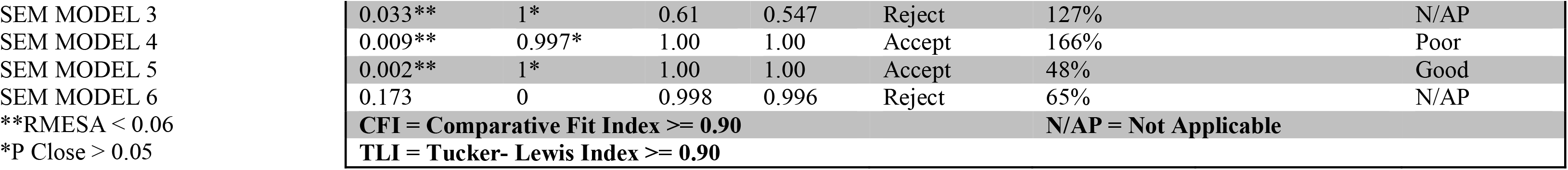
STRUCTURAL EQUATION MODEL.

## Results

The Freq. distributions of observed variables were displayed in Table No. 1f to 1g, 2a to 2b. Due to many number of SEM in this study, at first the model fit parameters [Table 7] were checked and final models were selected [Table 7]. According to obtained results SEM 1[Figure 1], SEM 2 [Figure 2], SEM 4 [Figure 4], SEM 5 [Figure 5] was accepted and SEM 3 [Supplementary Figure 3], SEM 6 [Supplementary Figure 6] was rejected.

**FIGURE:- 1.**
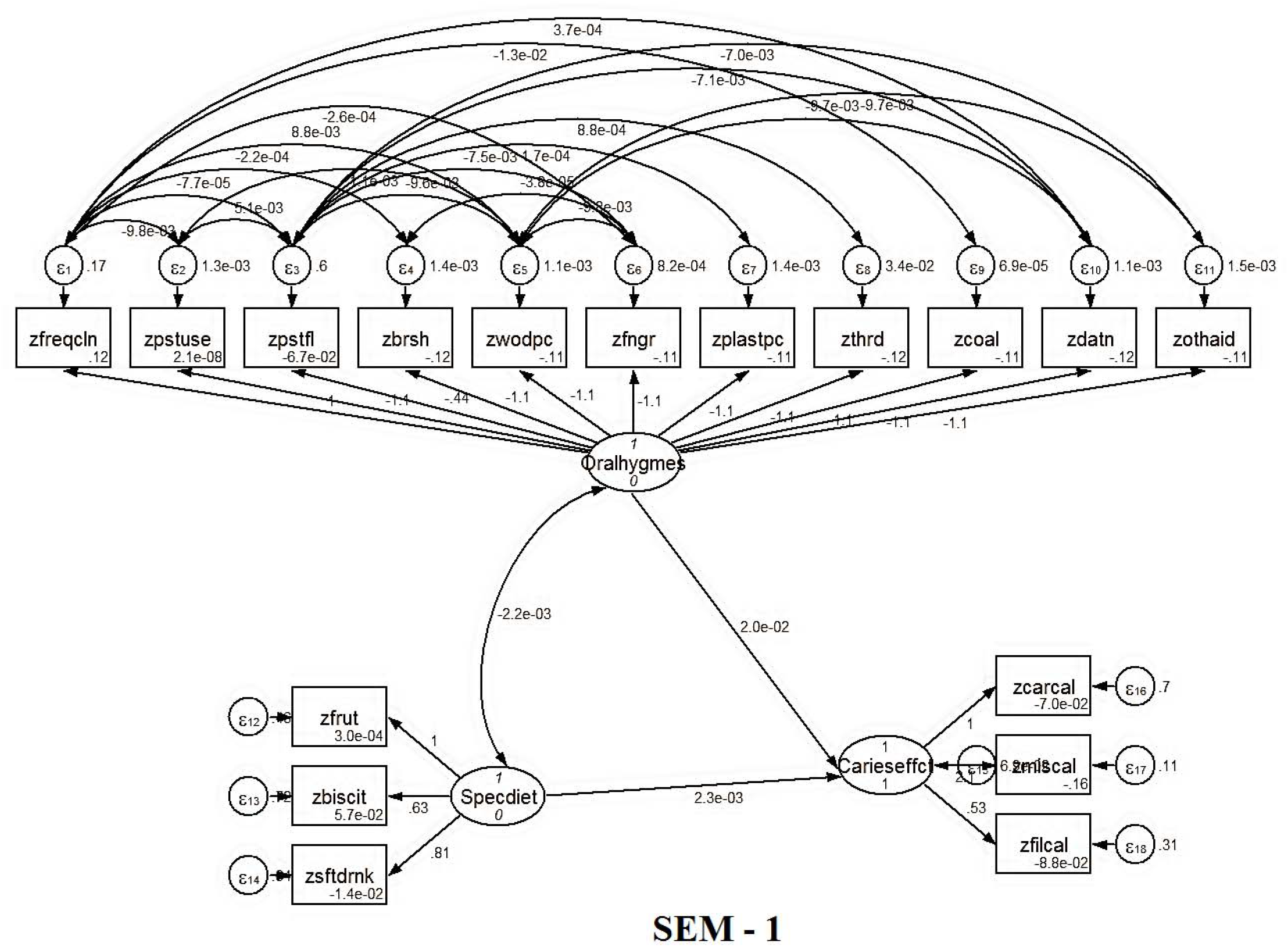
(In this model oral hygiene measure was positively associated with Caries effect [path coefficient (β) = 0.02, P < 0.05] and specific dietary habit was also positively associated with caries effect [path coefficient (β) = 0.0023, P < 0.05]. Oral hygiene measure was negatively co varying with specific dietary habit [-0.0022].)

**FIGURE:- 2.**
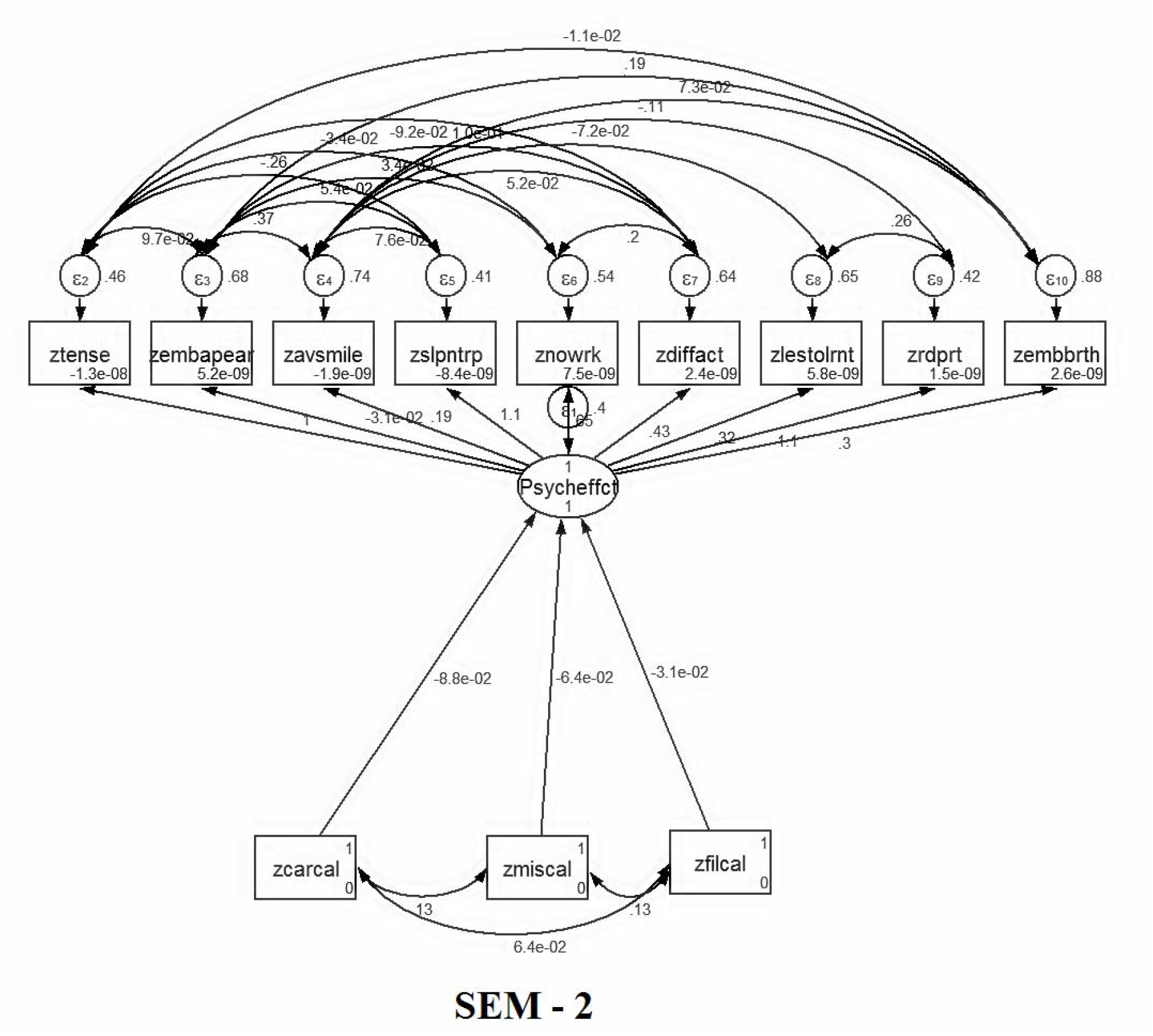
total number of carious teeth, total number of missing teeth and total number of filled teeth present in every subject was negatively associated with psychological effect [path coefficients (β) respectively −0.088,−0.064,−0.031, P< 0.05]. This thing proved that higher the dental problems less will be the psychological peace.

**FIGURE:- 4.**
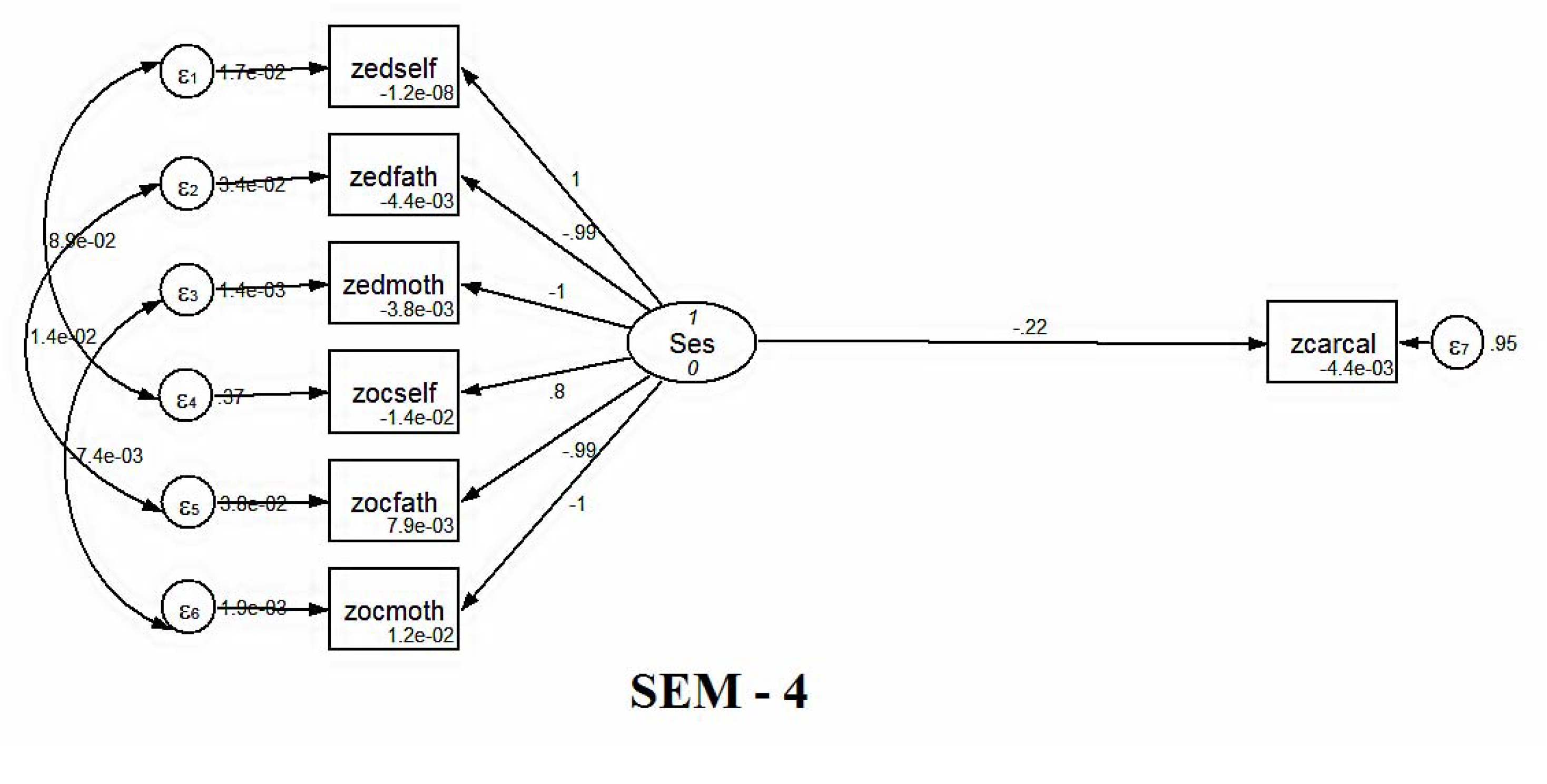
(In this model socio-economic status of the subjects were negatively related to total number of carious teeth present in each subject [path coefficient (β) = −0.22, P< 0.05].)

**FIGURE:- 5.**
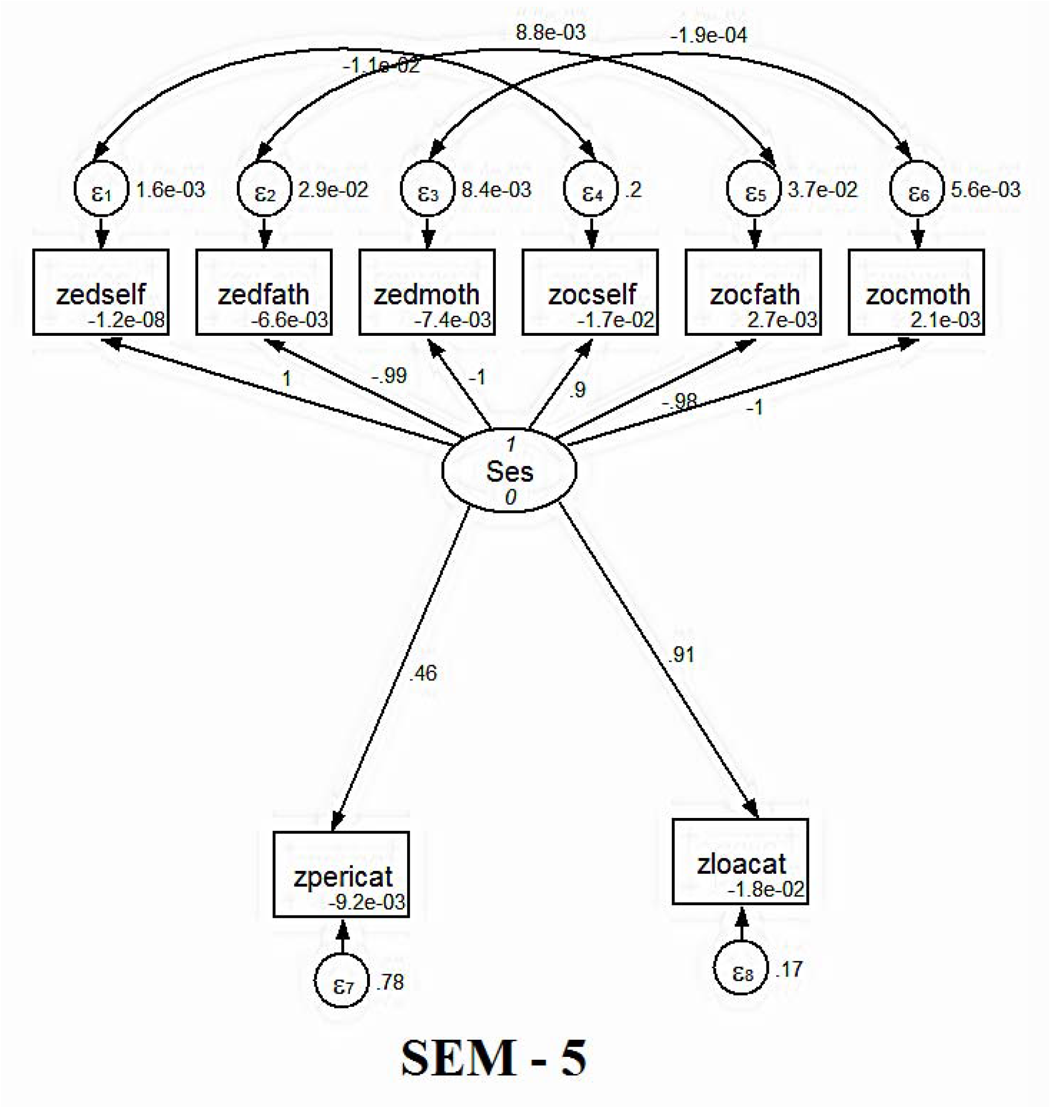
(In this model socio-economic status of the subjects were positively related to the poor periodontal condition [path coefficient (β) = 0.46, P< 0.05] and loss of periodontal attachment [path coefficient (β) = 0.91, P< 0.05].)

### SEM 1[Figure 1]:-

In this model oral hygiene measure was positively associated with Caries effect [path coefficient (β) = 0.02, P < 0.05] and specific dietary habit was also positively associated with caries effect [path coefficient (β) = 0.0023, P < 0.05]. Oral hygiene measure was negatively co varying with specific dietary habit [−0.0022]. That means bad oral hygiene measure (such as not using fluoridated tooth paste / using finger to clean teeth etc.) had a role to develop the carious condition of teeth. Eating sticky sweet food containing glucose or fructose, soft drinks etc. also had a positive role to develop dental caries. Caries effect was constructed by total carious teeth, total missing teeth due to caries, total filled teeth after caries removal present in subject’s oral cavity. So it caries effect was an indicator of total lifetime caries experience of the subject.

### SEM 2 [Figure 2]:-

In this model some exceptional characteristics were observed. RMSEA = 0, CFI = N/A because RMSEA of null model is 0.134774 which was less than 0.158. After calculation it was observed that χ^2^_k_/df_k_ < χ_0_^2^/df_0_ < 1[where χ^2^_k_/df_k_ for model =0.2749, χ_0_^2^/df_0_ for null model = 0.77212]. This thing proved that the baseline model was fitted very well with the data. As a result the TLI value = −2.182 was obtained. Total number of carious teeth, total number of missing teeth and total number of filled teeth present in every subject was negatively associated with psychological effect [path coefficients (β) respectively −0.088,−0.064,−0.031, P<0.05]. This thing proved that higher the dental problems less will be the psychological peace.

### SEM 4 [Figure 4]:-

In this model socio-economic status of the subjects were negatively related to total number of carious teeth present in each subject [path coefficient (β) = −0.22, P< 0.05]. So, poor socio-economic status had high influence in development of dental caries.

### SEM 5 [Figure 5]:-

In this model socio-economic status of the subjects were positively related to the poor periodontal condition [path coefficient (β) = 0.46, P< 0.05] and loss of periodontal attachment [path coefficient (β) = 0.91, P< 0.05]. After measurement obtained by CPI and LOA [according to WHO criteria], the CPI was further categorized as no periodontal problem, periodontal problem present and recording not required. The LOA was further categorized as no periodontal attachment loss, periodontal attachment loss present and recording not required. The basis of this categorization is mentioned before. All over it was proved that poor socio-economic condition was one of the reasons behind the periodontal disease development.

### Artificial Neural Network Model [ANN]:-

Artificial Neural Network [ANN] model with multilayer perceptron [MLP] were constructed by using same set of dependant and independent variables corresponding to each structural equation model [Table No. 4].

### ANN Model 1[Figure 7] and ANN Model 4 [Figure 13]:-

Both the model had continuous outcome/ dependant variable [Table 4]. ANN Model 1 and ANN Model 4 showed a large change of sum of square error from testing to training set of data, along with these average overall relative error (in %) was very high [Table 8]. COV of testing and training were also calculated as

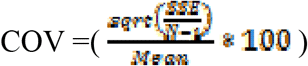

which showed 12% decrease in COV from training to testing. ANN Model 4 showed 16% increase in COV from training to testing [Table 9]. Average overall relative error was an indicator of inaccurate prediction done by the model.

**FIGURE:- 7.**
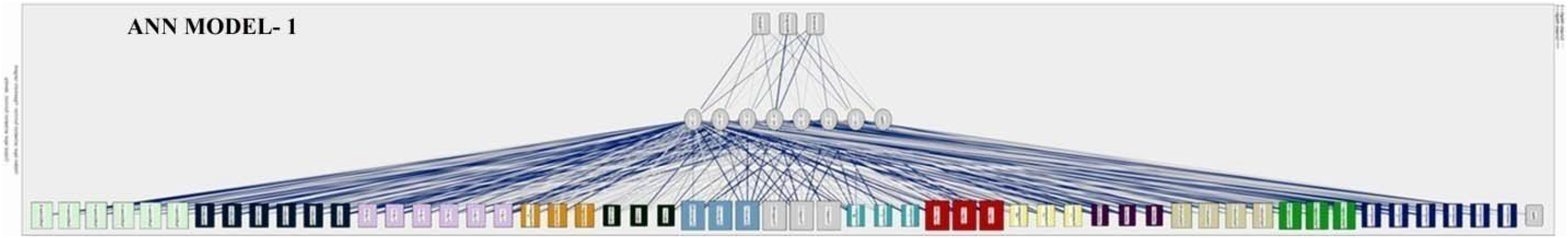
(ANN Model 1 showed the (%) of inaccurate prediction in training, testing, and holdout sample were respectively 96.8%, 97.7%, 98.8%.)

**Table No. 8:-.**
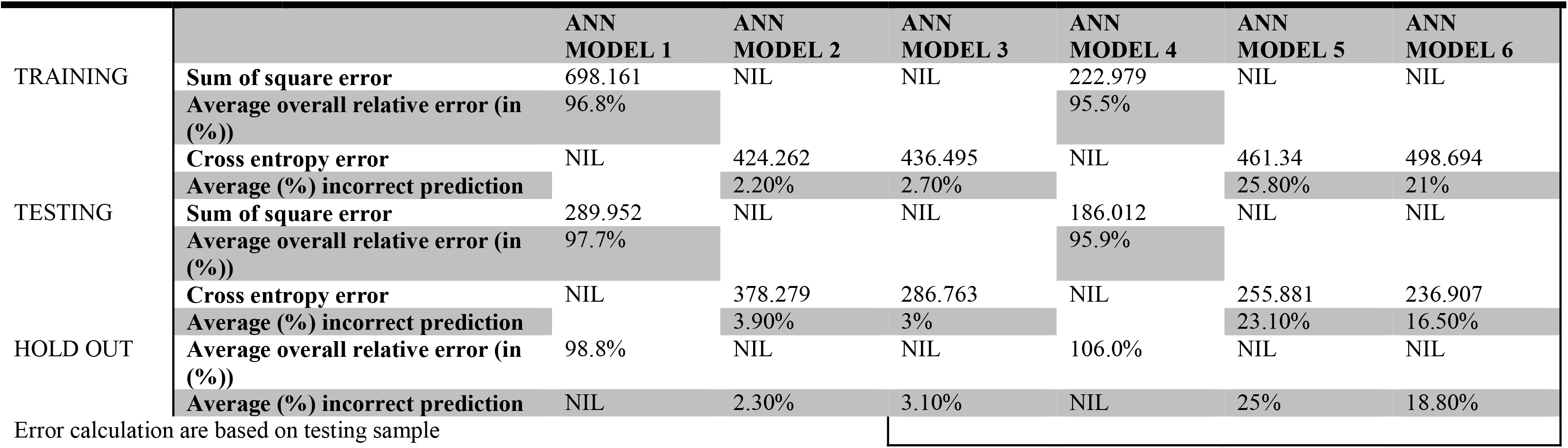
ARTIFICIAL NEURAL NETWORK MODEL SUMMARY.

**Table No. 9:-.** Covariance Calculation.

|  |  | ANN<br>MODEL 1 | ANN<br>MODEL 4 |
| --- | --- | --- | --- |
| TRAINING | COV | 183.2581435 | 64.29102768 |
| TESTING | COV | 160.9426695 | 74.7740775 |

ANN Model 1 showed the (%) of inaccurate prediction in training, testing, and holdout sample were respectively 96.8%, 97.7%, 98.8%. ANN Model 4 showed the (%) of inaccurate prediction in training, testing, and holdout sample were respectively 95.5%, 95.9%, 106.0%. So, in overall those ANN Models were unstable and gave huge amount of inaccurate prediction. As a result those ANN Models were rejected [Table 8].

### ANN Model 2 [Figure 9]:-

In this model cross entropy error was decreased 11% from training to testing sample. Average (%) of incorrect prediction in training, testing and hold out sample were respectively 2.2%, 3.9%, 2.3%. So this model was accepted [Table 8]. According to this model filled teeth present in oral cavity was the best predictor (had normalized importance of 100%) [Supplementary Figure 10].

**FIGURE:- 9.**
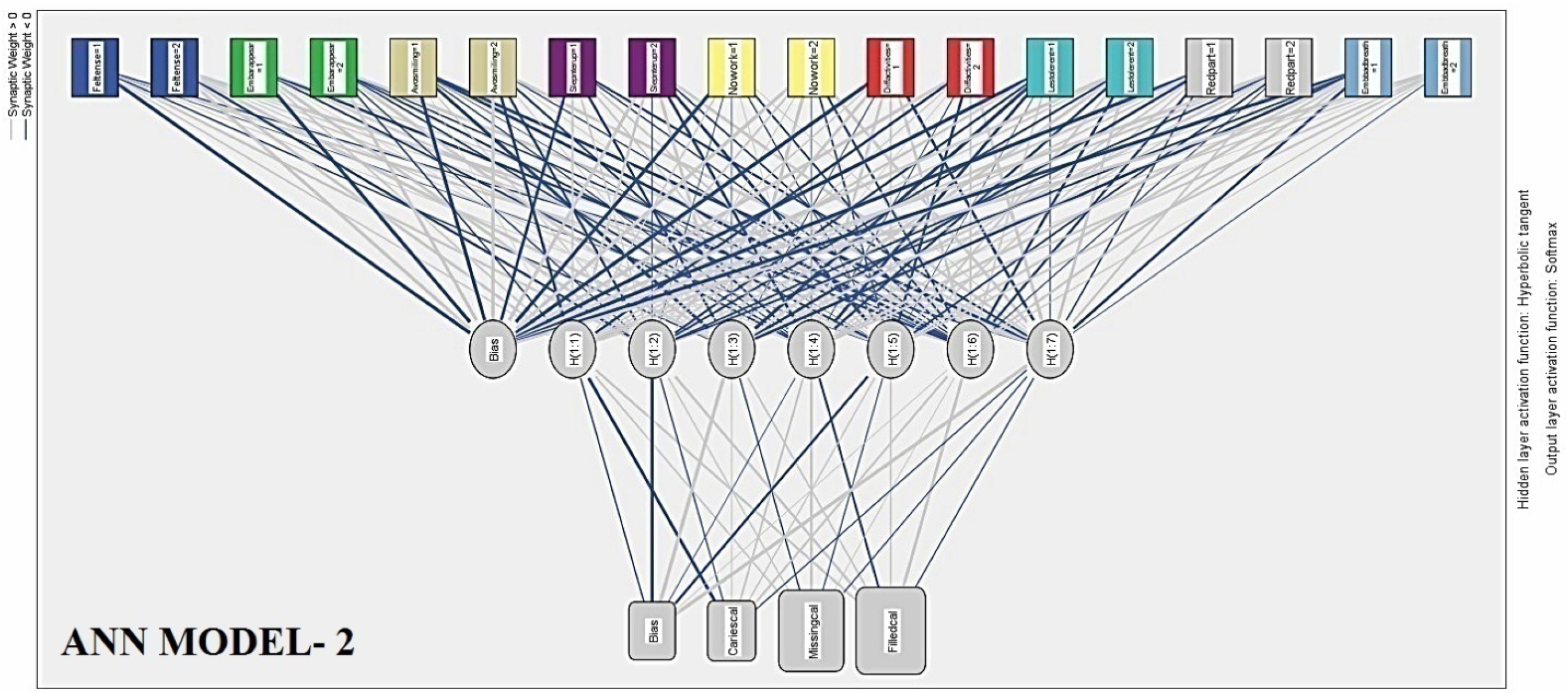
(Average (%) of incorrect prediction in training, testing and hold out sample were respectively 2.2%, 3.9%, 2.3 %.)

### ANN Model 3 [Figure 11]:-

In this model cross entropy error was decreased 34% from training to testing sample. Average (%) of incorrect prediction in training, testing and hold out sample were respectively 2.7%, 3.0%, 3.1%. So this model was accepted [Table 8]. According to this model missing teeth in oral cavity was the best predictor (had normalized importance of 100%) [Supplementary Figure 12].

**FIGURE:- 11.**
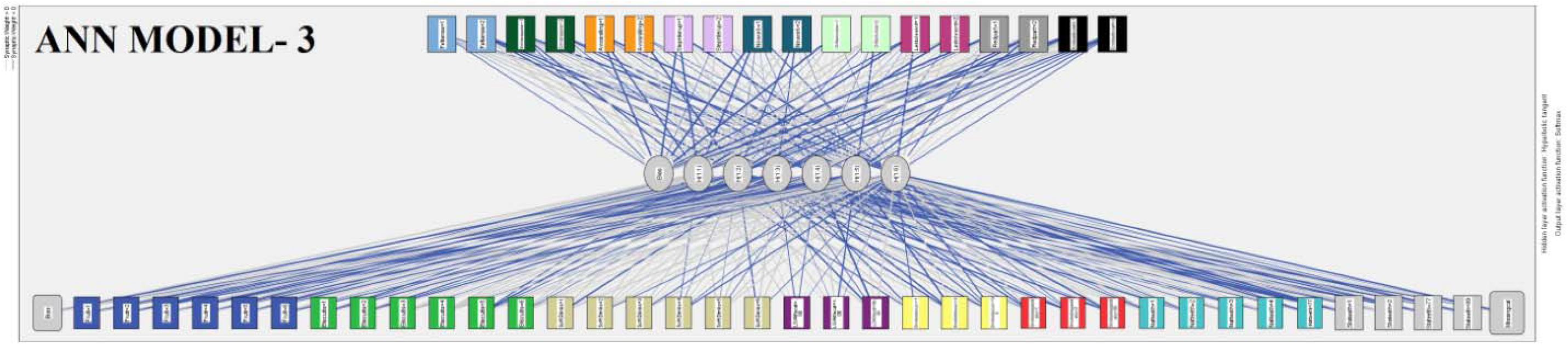
(Average (%) of incorrect prediction in training, testing and hold out sample were respectively 2.7%, 3.0%, 3.1%.)

**FIGURE:- 13.**
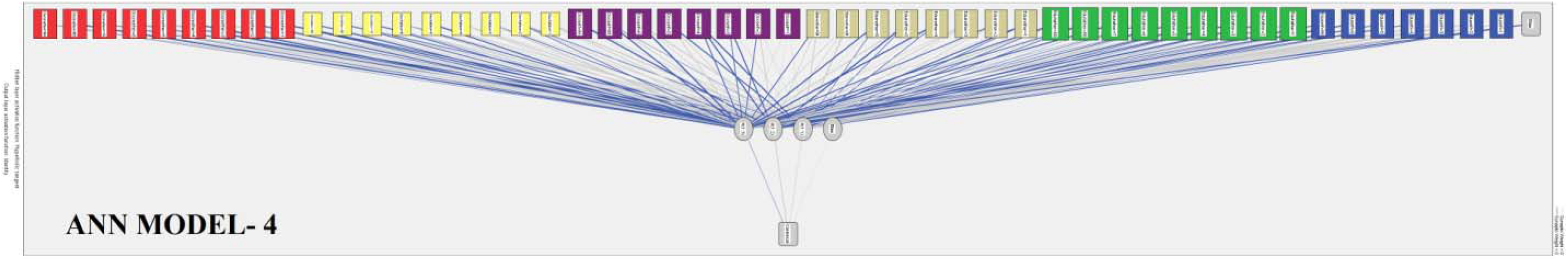
(ANN Model 4 showed the (%) of inaccurate prediction in training, testing, and holdout sample were respectively 95.5%, 95.9%, 106.0%.)

### ANN Model 5[Figure 15]:-

In this model cross entropy error was decreased 45% from training to testing sample. Average (%) of incorrect prediction in training, testing and hold out sample were respectively 25.08%, 23.10%, 25.0%. So this model was accepted [Table 8]. According to this model occupation of subject was the best predictor (had normalized importance of 100%) [Supplementary Figure 16].

**FIGURE:- 15.**
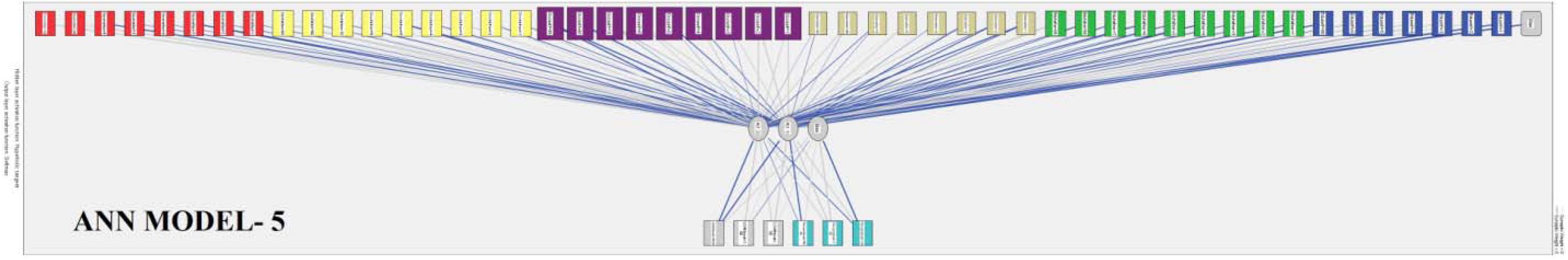
(Average (%) of incorrect prediction in training, testing and hold out sample were respectively 25.08%, 23.10%, 25.0%.)

### ANN Model 6 [Figure 17]:-

In this model cross entropy error was decreased 52% from training to testing sample. Average (%) of incorrect prediction in training, testing and hold out sample were respectively 21.0%, 16.50%, 18.8%. So this model was accepted [Table 8]. According to this model current smoking habit was the best predictor (had normalized importance of 100%) [Supplementary Figure 18].

**FIGURE:- 17.**
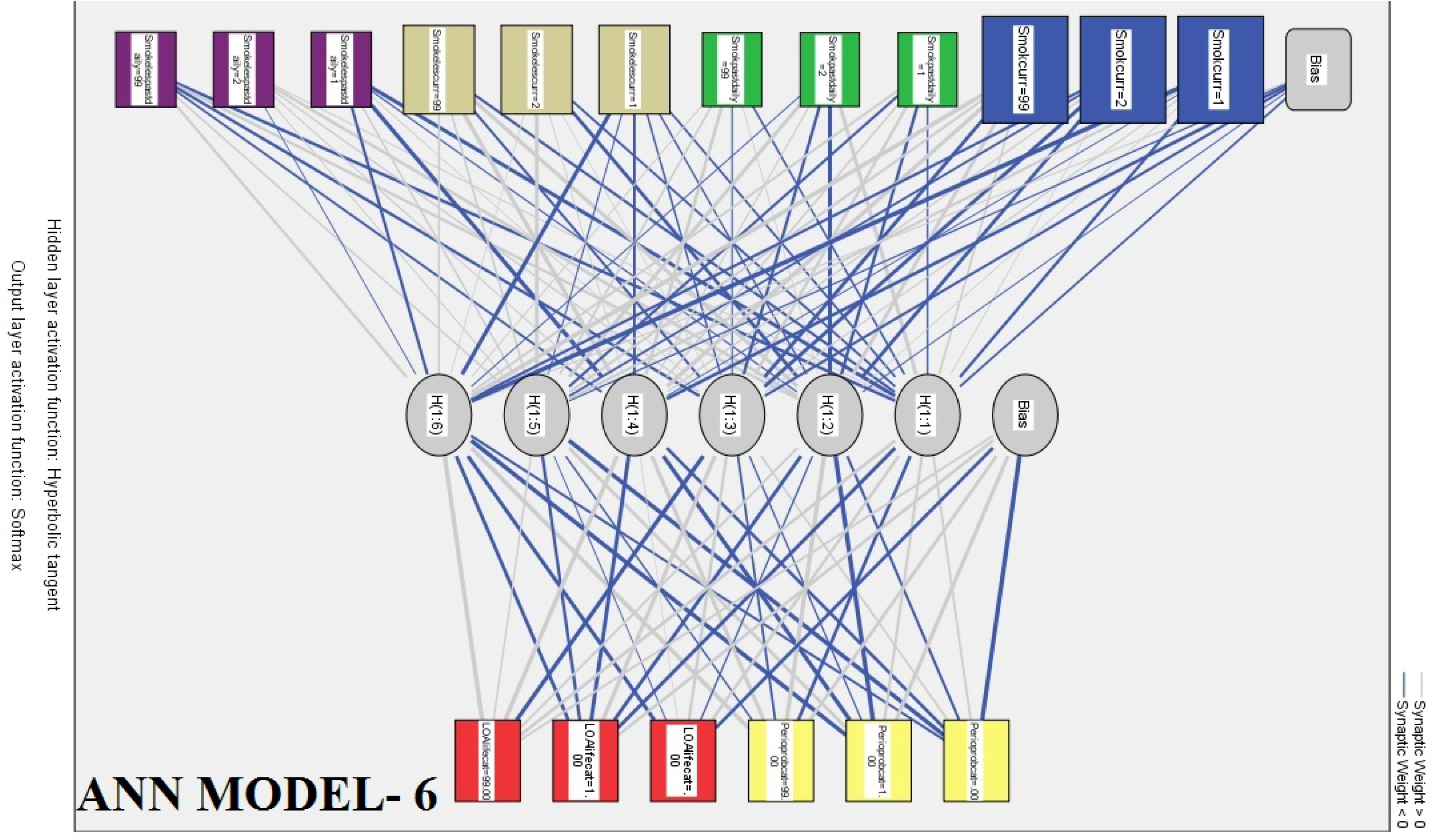
(Average (%) of incorrect prediction in training, testing and hold out sample were respectively 21.0%, 16.50%, 18.8 %.)

### Comparison of Artificial Neural Network Models and Structural Equation Models: -

At first predicted output was calculated from each six SEM. Later average overall relative error (in %) was calculated for each six SEM. Purpose behind this was to compare the SEMs to each other and with ANN Models in prospect of prediction capacity. The obtained values were presented in [Table No. 7 Final comment]. Which showed that SEM 5 predicted the dependant variable 52% correct (average overall relative error = 48%). SEM 1 predicted the dependant variable 18% correct (average overall relative error = 82%). In case of SEM 2 and SEM 4 the average overall relative error were 107% and 166% respectively, which proved that from the predictive point of view these two models performed very poor. SEM 3 and SEM 6 were not considerable in this predictive performance comparison because they were already rejected due to poor fit earlier [Table 7].

## Discussion

### Discussion on Statistical Models: -

Artificial Neural Network Models were suitable for predictive analysis purpose but it was not suitable to establish the cause and effect relationship. ANN was able to detect the most important and least important predictors but SEM was not able to do that. SEM produced some valid cause and effect relationship [22] among the observed variables which will help us to understand the exact structure and mathematical relationship among them [Table No. 10].

**Table No. 10:-.**
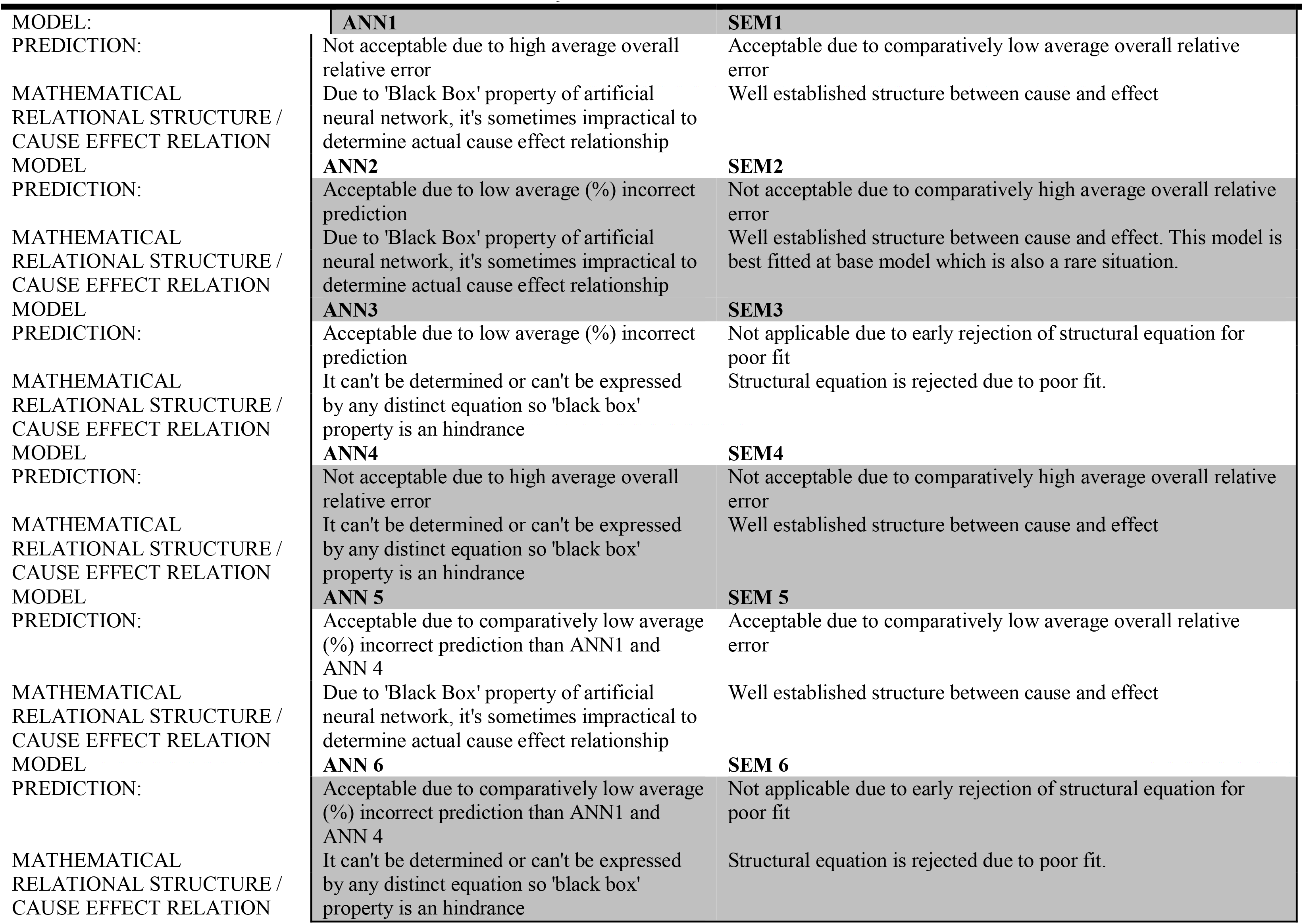
COMPARISON BETWEEN STRUCTURAL EQUATION MODEL AND ARTIFICIAL NEURAL NETWORK MODEL.

### Discussion on clinical relationship: -

Results shows that study’s hypotheses are proved correct in case of SEM 1, 2, 4 and 5. Oral hygiene habit measure of the population and their dietary habit are directly influencing the development of dental caries. Carious teeth, missing and filled teeth are affecting the mental health of the persons. Socio-economic condition of a person is directly involved in development of dental caries and periodontal disease. The KABP Model is composed of Knowledge (K), Attitude (A), Belief (B), and Practice (P) but in reality belief is very similar to attitude; therefore the KABP model is more specifically KAP[14] is always applied in oral health promotion, through oral health education[15,16]. People’s ‘Knowledge’ and ‘Attitude’ about oral health are the main reason behind their ‘Practice’ [7].

Every study have some strength and limitations, so strength of this study are:- First the SEM method is more superior than multiple regression modeling which can only the direct effects, no latent factor cannot be included in multiple regression modeling. Second a large random sample that contained a variety of socio-economic groups and age-gender groups were used in this study so sample is the perfect reflection of general population, which will increase the external validity of results. Third prediction was of outcome and detection of important predictor variable by ANN. In future more importance will be given to these relationships and predictor variable for formulation of health policy.

#### Limitations of the study are: -

First dental caries, periodontal disease and psychological problems are caused by multiple different factors [13] and studies on the roles of genetics, biology, social environment, mental health, physical environment, health influencing behaviors and medical care are critical for a complete understanding of their influences on oral and mental health; the present study merely explained some of those influences. Second the development of dental caries, periodontal disease and psychological problems are chronic and progressive in nature. As it is a cross sectional study to establish the relationship among the factors, there is a need to conduct longitudinal study for better understanding.

## Conclusion

In conclusion findings obtained from this study have important application for policy making with regard to public health and people should be made to realize that proper diet and oral health maintenance are keys to get a disease free oral condition. Caring their teeth and gums will promote a healthy life style and also help to reduce the psychological disturbances caused due to oral diseases. Additionally public education programs should be advocated more broadly and oral health care service should be improved in socio-economically deprived areas.

## Abbreviations

Refer to the List of abbreviations table.

## Competing interests

The author declares that no potential conflict Of interest with respect to authorship and/or publication of this article.

## Author’s contribution

Author was partly involved in study design, methodology development, and questionnaire preparation. Completely involved in clinical examination, field work, statistical analysis and article preparation.

## Acknowledgements

This study was funded by WHO [SEARO], New Delhi, India (on year of 2014). The study was conducted by Centre for Dental Education and Research [AIIMS], New Delhi, India. And thanks should be given to the following persons and institutions: Prof. Naseem Shah [former chief CDER, AIIMS, New Delhi, India], Dr. Vijay Prakash Mathur [Additional Professor, CDER, AIIMS, New Delhi, India], Mr. Prabhat Sangal [Assistant Professor, School of Sciences, IGNOU, New Delhi, India], Dr. Saradindu Ghosh [Qa&Qc Manager, Patanjali Pvt. Ltd, Haridwar, India] and whole study participants.

## Author detail

Former Senior Research Fellow at Centre for Dental Education and Research, New Delhi, India.

---

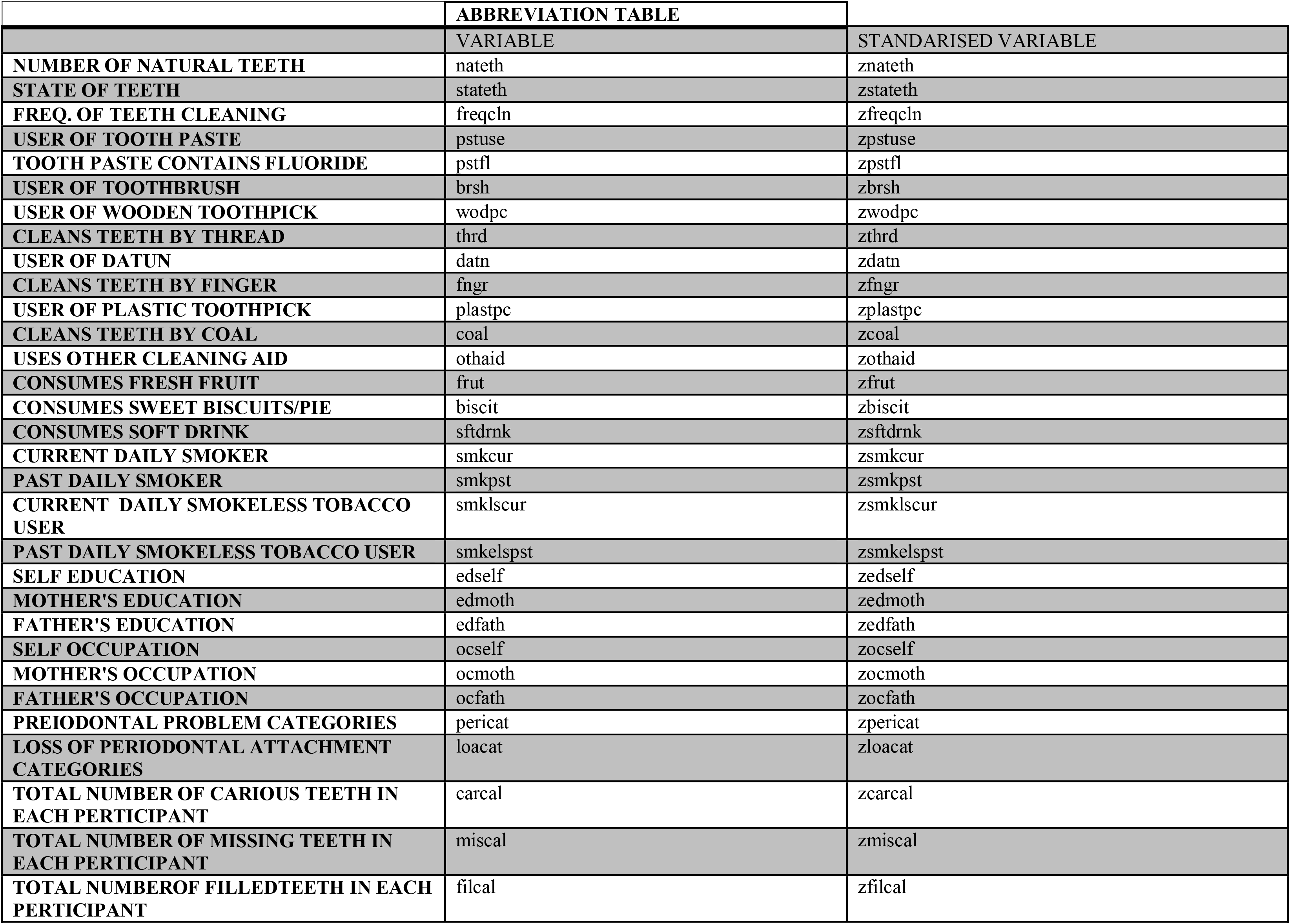

